# *Pichia pastoris* growth - coupled heme biosynthesis analysis using metabolic modelling

**DOI:** 10.1101/2023.05.13.540629

**Authors:** Agris Pentjuss, Emils Bolmanis, Anastasija Suleiko, Elina Didrihsone, Arturs Suleiko, Konstantins Dubencovs, Janis Liepins, Andris Kazaks, Juris Vanags

## Abstract

Soy legHemoglobin is one of the most important and key ingredients in plant-based meat substitutes that can imitate the colour and flavour of the meat. In order to improve the high-yield production of legHemoglobin protein and its main component - heme in the yeast *Pichia pastoris*, glycerol and methanol cultivation conditions were studied. Additionally, *in-silico* metabolic modelling analysis of growth-coupled enzyme quantity, suggests metabolic gene up/down-regulation strategies for heme production. First, cultivations and metabolic modelling analysis of *P. pastoris* were performed on glycerol and methanol in different growth media. Glycerol cultivation uptake and production rates can be increased by 50 % according to metabolic modelling results, but methanol cultivation – is near the theoretical maximum. Growth-coupled metabolic optimisation results revealed the best feasible upregulation (33 reactions) (1.47 % of total reactions) and 67 downregulation/deletion (2.98 % of total) reaction suggestions. Finally, we describe reaction regulation suggestions with the highest potential to increase heme production yields.

## Introduction

Meat is one of the significant sources of dietary protein. It is frequently recognised as a high-quality protein source due to its nutritional qualities and favourable sensory properties such as texture and flavour. However, the rising global population has led to a rise in the production and consumption of meat around the world^1^, which in turn has raised environmental concerns regarding the usage of land and water, as well as its impact on pollution and climate change, greenhouse gas emissions, and the loss of biodiversity^2^. Recommendations to limit meat consumption have been made due to the fact that meat is a critical source of protein and in numerous high-income countries protein utilisation outperforms dietary necessities^3^. Plant-based meat can be an efficient alternative to livestock, as for the production of the primary plant proteins, fats, carbohydrates and vitamins are used. Plant-based meat is defined as a meat-like substance made from vegetarian-friendly ingredients like proteins (from soy and potatoes), fat (from coconut and sunflower oils), carbohydrates (potato starch, corn starch), nutrition additives (yeast extract, vitamins) and flavours (beet juice extract, apple extract and plant haemoglobin)^4^. The market for plant-based meat is anticipated to reach a significant milestone of $30.9 billion by 2026^5^ due to the rise in the number of vegetarians, vegans, and flexitarians in Europe during the past several years^6^. The switch from an animal to a non-animal diet is also influenced by religious considerations and high production costs^7^.

Heme-containing proteins are considered as one of the crucial flavour and colorant contributors in animal red meat^8^. Also, hemoprotein addition to artificial meat or its analogues has been proven to display great potential to mimic the taste and smell similar to that of traditional meat. LegHemoglobin (legH) is approved by the United States Food and Drug Administration (FDA) as a food colouring additive and can be used equivalently to haemoglobin and myoglobin in both vegetarian and non-vegetarian diets^9^.

LegHemoglobins, are present in the root nodules of leguminous plants. In plant-based meats legH performs a parallel role: it unfolds upon cooking, releasing its heme cofactor to catalyse reactions that can transform the same biomolecules, isolated from plant-based sources, into the array of compounds that comprise the unique flavour and aroma of traditional meat^10^. The use of legH protein as a food additive also circumvents the contradictions of animal material used in the diet, especially for vegetarians.

Many attempts in biotechnology have already been made to increase hemeprotein yields, thus ensuring feasibility of such technology for industrial scale production. For example, to improve heme-containing protein production the addition of 5-aminolevulinic acid (ALA) to the growth medium and overexpression of heme transporter genes^11,12^ was studied, furthermore, other groups studied extracellular hemin addition, which provided only 5 - 10 % increase of heme-containing protein mass in respect to the total protein content^13^. In heme-containing protein production, extracellular precursor addition seems to not be an economically feasible approach and further studies should be focused to achieve commercially viable intracellular hemoglobin, myoglobin and legH production through genetic engineering.

Currently, microorganism platforms are the most perspective way of manufacturing LegH in commercial amounts, for further integration into plant-based meat. Several legH-producing recombinant microorganisms like: *Candida spp*., *Hansenula spp*., *Pichia spp*., or *Toruplosis spp*. representatives, which are methylotrophic yeast and can utilize alcohols, also methanol, have been previously used for accomplishing the mentioned tasks^14^. Furthermore, alternative microorganisms, like *Kluyveromyces lactis*, which are capable of utilising cheap dairy industry by-products, like cheese whey, have been studied^15^. Another study reported that it was possible to increase legH secretion by 83-fold in a genetically engineered *P. pastoris* in respect to the wild-type strain, by manually overexpressing globin and heme pathways genes separately^16^. But a more recent study showed that by combining systems biology and genetic engineering approaches, heme production was increased 70-fold in *Saccharomyces cerevisiae*. This was done using metabolic modelling coupled with growth heme production analysis on the genome level, where the most promising analysis results were genetically manipulated and experimentally tested^17^.

In the last decade, genome-scale metabolic modelling is increasingly being used as an auxiliary tool for facilitating various genetic engineering tasks and for predicting potential effects. Metabolic models effectively encode gene-protein-reaction (GPR) associations^18^ and explain genotype-phenotype interaction in unicellular and multi-cellular organisms^19,20^.

Metabolic modelling allows to predict the organisms phenotypes biotechnological properties and responses^21^, predicts environmental perturbation impacts on the metabolism and potential carbon distribution^22,23^. Genome-scale metabolic model (GEM) application in biotechnology already has proven its importance and helped to develop new nutrient uptake innovations^24^, novel software developments^25,26^, created new biotechnology applications^27,28^.

GEM investigation has guided the development of strains with optimised yields of industrial metabolites (e.g., sesquiterpenes, vanillin, bioethanol, 2,3-butanediol, succinate, amorphadiene, β-farnesene, and dihydroxyacetone phosphate, 3-hydroxy propionate, fumaric acid)^29,30^.

One of the most common approaches to metabolic engineering and computational biology during the last decade has spread to link the production of the desired metabolite with coupled growth^31^. This method is very effective in new strain design because metabolic production becomes a necessary by-product for cell growth. Furthermore, to increase desired metabolite production yields and production capabilities it is possible adapting laboratory evolution by selecting for maximum growth^32,33^.

Over the past 20-30 years, the *P. pastoris* expression system has been extensively used for the production of various recombinant proteins for both research and industrial applications. This methylotrophic yeast is highly suitable for foreign protein expression due to several key features, including ease of genetic manipulation, high-frequency DNA transformation, cloning by functional complementation, high levels of intracellular and extracellular protein expression, and the ability to perform higher eukaryotic protein modifications (such as glycosylation, disulfide bond formation, and proteolytic processing)^34,35^. In addition, relatively low levels of native secreted proteins allow for simple purification of the secreted recombinant proteins. Taking into account the economic factors (such as high cell growth in minimal media and high product stability in prolonged processes) and the advanced genetic techniques available, *P. pastoris* is clearly the system of choice for heterologous protein expression^36,37^. The versatility of *P. pastoris* is reinforced by the high yields of various recombinant proteins achieved in this microorganism^16,38–40^.

Experimental cultivations on glycerol and methanol for legH protein production allowed us to find out that there exists 50 % more potential maximal theoretical *P. pastoris* biotechnological potential for fast biomass increase in bioreactors, but legH protein production compared with GSM data showed that external substrate addition is close to maximal bioproduction potential. Therefore, it indicated a possibility that to increase the production of legH protein in economically viable quantities, intracellular reactions/gene manipulations must be carried out. This study was aimed to determine a list of the potential reaction manipulation strategy for high heme biosynthesis through systematic growth coupled analysis in *P. pastoris* genome-scale metabolic model.

First, we developed an algorithm for growth-coupled enzymatic reaction and corresponding gene up-down regulation impact analysis on heme production. Then the algorithm was applied to *P. pastoris* GSM and we found that only 500 reactions from 2243 and the most promising reactions for downregulation/deletion were 33 reactions and directly proportional 67 reactions.

For upregulation, the algorithm suggests that heme-BIOSYNTHESIS-II and PWY-5189 are the best candidates and formaldehyde assimilation III (dihydroxyacetone cycle) (P185-PWY) is the less effective candidate. For downregulations/deletion suggestions, most potential suggestions are different amino acid metabolism, where they possibly compete with heme biosynthesis for glycine.

## Results and Discussion

To develop *P. pastoris* reactions regulation design sets for genetic engineering purposes and increase intracellular heme production rate, we chose to use published *P. pastoris* iMT1026 GEM, which consists of 1708 metabolites, 2243 reactions, 2 different biomass compositions for glycerol and methanol fermentation optimisations, has 9 different compartments and transport intercompartment^41^. iMT1023 model was updated and missing mandatory information filled to be compatible with CobraToolbox 3.0 ^44^.

GSM includes metabolite name, charged formula and compartment name in the cell. As iMT1026 has no metabolite or reaction database references, thus it was difficult to test new functionality in comparison with scientific literature. GSM is considered to work at pH 7.2, 32 °C environment temperature. To test GSM’s possibility to correctly simulate physiologically relevant conditions and analyse heme biosynthesis growth-coupled production, we performed experimental glycerol and methanol cultivations for legH production. We assumed that the legH production yields are approximately the same as heme intracellular production yields in methanol cultivation experiments.

*P. pastoris* for heme production using a diauxic cultivation process, where the preferred substrate is glycerol and after glycerol concentration decreases, the growth of methanol is induced. Methanol is used to induce recombinant protein production^45^. The same approach was exploited for heme production. When comparing *P. pastoris* cultivation on both substrates the measured specific growth rate (**μ**) utilizing glycerol was 6.3 - 4.75-fold higher (∼ 0.19 h^-1^) than consuming methanol (0.04 - 0.03 h^-1^), which is in the scope of previously published data (1.7–8.5 times faster)^45^. The O_2_ consumption rate on glycerol growth was 2.17 - 2.29 mmol*g^-1^ * h^-1^, but CO_2_ production was a little less (1.61 - 1.7 mmol*g^-1^ * h^-1^) than O_2_ consumption (Supplementary materials 1).

After obtaining growth, O_2_ and CO_2_ uptake rates using *Optek* and *Incyte* driven glycerol and methanol cultivations the GSM model was accordingly constrained with experimentally measured exo-metabolomics data. After *P. pastoris* GSM optimisations, results suggested that the GSM model can achieve a steady state and produce heme. Comparing the GSM optimisation results with experimental data, we concluded that experimental results driven by Optek are a little higher than experiments driven by Incyte, but in general have the same conclusions. Comparing the glycerol and methanol growth conditions, the first experimental CO_2_ production (2.75-2.9-fold) and O_2_ consumption (2.4-2.5-fold) is lower than the model suggested predictions, which prompts the conclusion that in glycerol growth conditions there exists potential to increase metabolic rates. Although in methanol growth conditions we can see that CO_2_ production (0.04-0.3-fold) and O_2_ consumption (0.14-0.41-fold) is less low than GSM predictions and relatively close to the maximal heme production. The same conclusions have been made by earlier studies^11–13^, which concluded that extracellular addition of substrates is a less effective strategy and has its own limitations, than genetic engineering potential, like *S. cerevisiae* GSM driven 70-fold increase of intracellular heme production^17^.

Thus, next step in this study was to use *P. pastoris* GSM to make a systemic reaction impact analysis on potential heme production. Previously described experimental and GSM optimisation results comparisons suggested a small potential to increase heme production. This could be one of the GSM modelling bottlenecks in the biomass reaction itself. In GSM exist biomass objective function^46^, which defines necessary metabolic resources for cell process machinery and proliferation needs. But the biomass objective function data is experimentally measured in strict cultivation conditions, is expensive to measure and sometimes even has a large error distribution. Also, it is known that biomass objective function composition and macromolecule weighting factors can change during different environmental conditions^47^

In this study, we improved the previously published growth-coupled GEM algorithm and adjusted it to new GEM requirements. To exclude biomass objective function limitations, we decided that growth-coupled GEM algorithm functionality (see materials and methods) will systematically analyse all metabolic reactions’ impact on heme production. As a result, we found the most promising upregulation (33 reactions) (1.47% of total reactions) (Supplementary Materials 3 Table S3.7) and 66 downregulation/deletion (2.98 % of total) reaction candidates (Supplementary Materials 3 Table S3.8).

### Upregulation calculations

To increase heme intracellular production algorithm has suggested upregulating reactions from the dihydroxyacetone cycle combined with glycolysis reactions (P185-PWY from MetaCyc) and heme biosynthesis pathways: tetrapyrrole biosynthesis II (PWY-5189 from MetaCyc) and aerobic heme b biosynthesis I (heme-BIOSYNTHESIS-II from MetaCyc).

Interestingly the highest step-weighted factor for upregulation is mitochondrial pyrroline-5-carboxylate reductase. The conversion of glutamate to proline and vice versa is thought to play a role in ATP increase^48,49^. This could be related to 5-aminolevulinate synthase (ALASm), where Succinyl CoA (succoa) is consumed in large amounts as a 5-Aminolevulinate (5aop) precursor. In the mitochondrial TCA cycle Succinate--CoA ligase (ADP-forming) reaction, which is one of the suggested upregulation reactions is used to produce Succinyl CoA (SucCoA) and consume large amounts of ATP, which could be a limiting step for improved intracellular heme production if less ATP is available in mitochondria.

To increase intracellular heme bioproduction the most perspective upregulation reactions are 5-aminolevulinate synthase (ALASm), porphobilinogen synthase (PPBNGS), hydroxymethylbilane synthase (HMBS), uroporphyrinogen-III synthase (UPP3S) from heme b biosynthesis I pathway (heme-BIOSYNTHESIS-II) and uroporphyrinogen decarboxylase (UPPDC1), coproporphyrinogenase (CPPPGO), protoporphyrinogen oxidase (PPPGOm), protoporphyrin ferrochelatase (FCLTm) from tetrapyrrole biosynthesis II (from glycine) pathway (PWY-5189), which together forms heme biosynthesis metabolic pathway (Porphyrin and Chlorophyll Metabolism in GSM). These reactions have one of the highest step-weighted factors and have the highest impact on heme production increase as a whole pathway. In *S. cerevisiae* the HEM3 gene (in GSM HMBS reaction) was identified as rate limiting^50^, but recent studies in *P. pastoris* showed that HEM1 (in GSM ALASm)^16^ they analysed single gene upregulation of heme-BIOSYNTHESIS-II and PWY-5189 pathways and concluded that ALASm reaction upregulation had the best increase in legH production compared to other heme biosynthesis pathway reactions. The same conclusions indirectly suggest GSM pyrroline-5-carboxylate reductase step-weighted factor algorithm results. Thus, different results are found in different yeast species, which suggests heme production in various species is differently regulated. More the study suggested that HEM1 (ALASm), HEM2 (PPBNGS), HEM3 (HMBS) and HEM12 (FCLTm) upregulation showed a 41 % increase and all heme production genes overexpression led to 1.5-fold increased heme production, leaving the potential for much more improvements as it was recently found in *S. cerevisiae*^17^.

The next reaction group with step-weighted factor (11-14 %) are (a) formyltetrahydrofolate dehydrogenase (FTHFDH), methylenetetrahydrofolate dehydrogenase (NADP) (MTHFD), glycine hydroxymethyltransferase (GHMT2r), which are one-carbon metabolism pathway reactions responsible for the interconversion of different tetrahydrofolate (THF) forms. (b) Phosphoglycerate dehydrogenase (PGCD), phosphoserine transaminase (PSERT), phosphoserine phosphatase (L-serine) (PSP_L), glutamate dehydrogenase (NADP) (GLUDyi) and pyruvate carboxylase (PC), which in GSM are glycine (gly) precursors. The glycine is the precursor of heme-BIOSYNTHESIS-II and PWY-5189 pathways which produces heme.

As the last group of upregulation suggestions are the Pentose Phosphate pathway Dihydroxyacetone synthase (DAS) and Dihydroxyacetone kinase (DHAKx) from the methanol metabolism pathway, which together form formaldehyde assimilation III (dihydroxyacetone cycle) (P185-PWY). These reactions have less step-weighted factor (5-9 %) and have less impact on heme biosynthesis. These reactions are closely interconnected and allow methylotrophs to metabolise methanol to formaldehyde (fald), which later forms Glyceraldehyde 3-phosphate (G3P) and dihydroxyacetone (DHA) - the building block chemicals for all other metabolic processes, including of heme biosynthesis.

### Downregulation/deletions calculations

Relying only on an upregulation strategy to increase heme intracellular production is not sufficient enough. Previous studies on *Escherichia coli*^51^ proved that metabolic modelling proposed unnecessary metabolic process deletion for glycerol uptake allowed to increase succinate production yields. This demonstrated that some metabolic processes might be not essential and even lower product yield.

The corresponding reaction intracellular deletion can increase product yields not changing substrate consumption amounts. In this study, we found 66 individual reactions, in which flux rates change inversely proportional to heme production.

The largest group of downregulation/deletion candidates are different amino acids and their intermediate interconversion-related metabolism reactions with step-weighted factor 17.3. The exception is Aspartate 1-decarboxylase (ASP1DC) with a lower step-weighted factor 15. Group consists of 26 reactions from 66 (∼41 % of the total), involving 8 different amino acid pathways (Fig. 1) (Supplementary materials 3 Table S3.9).

**Figure 1.**
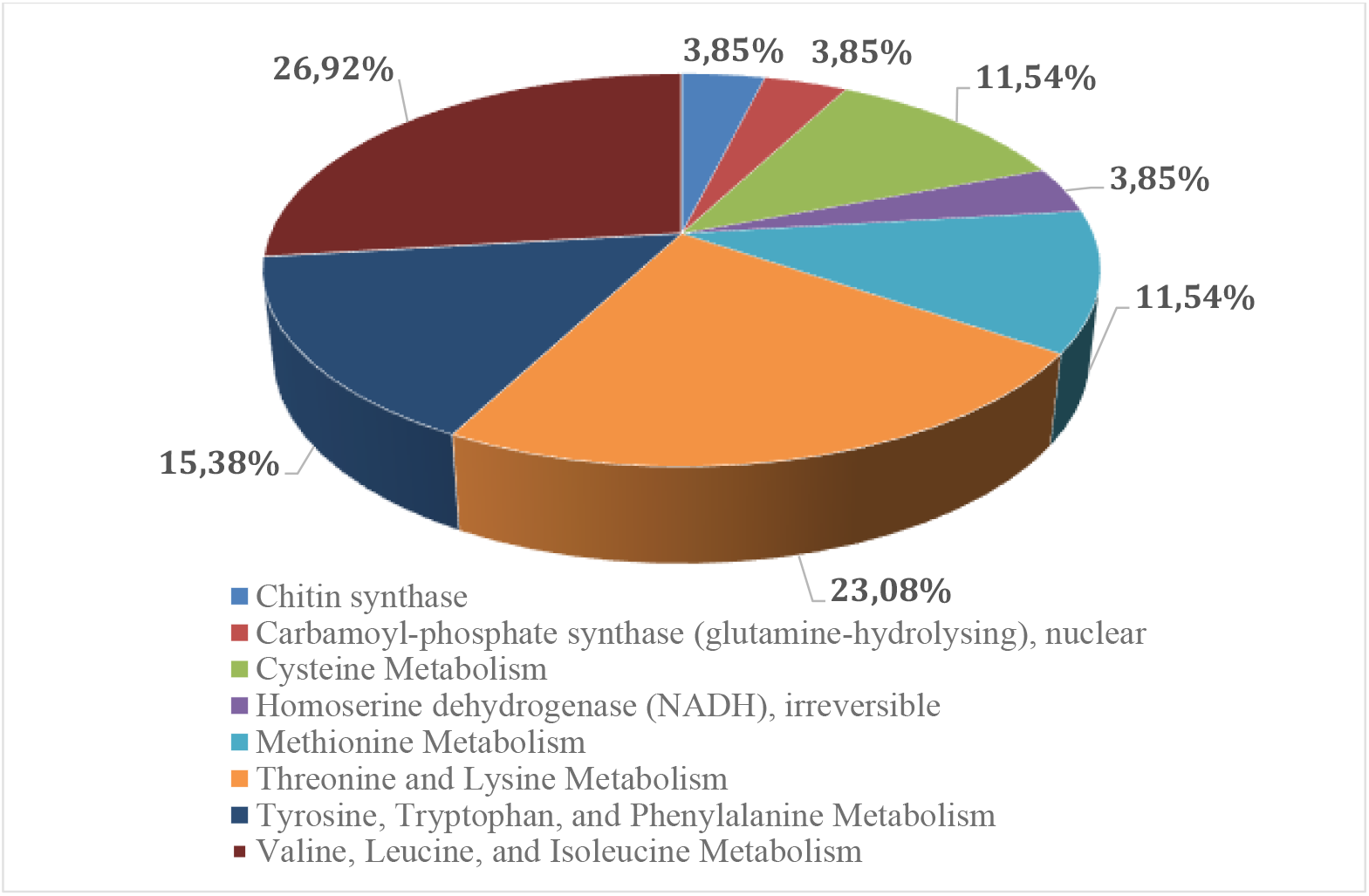
Fractional distribution of amino acids downregulation/deletion reaction candidates.

The largest amount of candidate reactions is:

- Valine, Leucine, and Isoleucine metabolic pathway: 7 reactions (25.93 %);
- Threonine and Lysine pathway: 6 reactions (22.22 %)
- Tyrosine, Tryptophan and Phenylalanine pathway: 5 reactions (18.52 %);
- Cysteine pathway: 3 reactions (11 %);
- Methionine pathway: 3 reactions (11 %)

We assume that in our experimental model, P. *pastoris* amino acid metabolism for *biomass growth* competes with heme biosynthesis for the glycine. Additionally, *P. pastoris* upregulate the C1 metabolism pathway in mitochondria to increase glycine synthesis necessary for heme (heme-BIOSYNTHESIS-II and PWY-5189) biosynthesis (see chapter “upregulations”)

The downregulation of amino acid metabolism could be related to the allocation of resources towards glycine and heme synthesis. Threonine is starting point for leucine and isoleucine synthesis, it is produced by the threonine aldolase from glycine. By lowering leucine, valine and isoleucine metabolism - the system retains more glycine to allocate for heme synthesis.

The second largest group is the Nucleotide Metabolism pathway 11 (∼16.5 % of total) reactions, in which the step-weighted factor is 17.3-18.1 %. The algorithm suggests that these reactions group downregulation/deletions are not essential and will not affect cell growth rate. Purine biosynthesis synthesis depends on glycine and folate supply - one glycine and two formyltetrahydrofolate molecules are necessary to yield purine molecules. Our results reveal, that DHFRim (dihydrofolate reductase, mitochondrial) should be downregulated to achieve max heme synthesis. There are 2 reaction chains from a 5,10-methylenetetrahydrofolate to tetrahydrofolate in yeasts and DHFRim is glycine dependent, while the parallel reaction chain is not.

Therefore, by keeping upregulating C1 metabolism and glycine synthesis and downregulating their consumption elsewhere - heme yield can be improved. Moreover, purine synthesis uses 4 ATP per 1 purine molecule; if down-regulating the purine synthesis pathway, some ATP could be reallocated to heme synthesis.

The third large group consists of 23 reactions from different metabolic pathways (Supplementary materials 3 Table S3.9), which the GSM algorithm suggests as potential downregulation/deletion reactions (Fig. 2).

**Figure 2.**
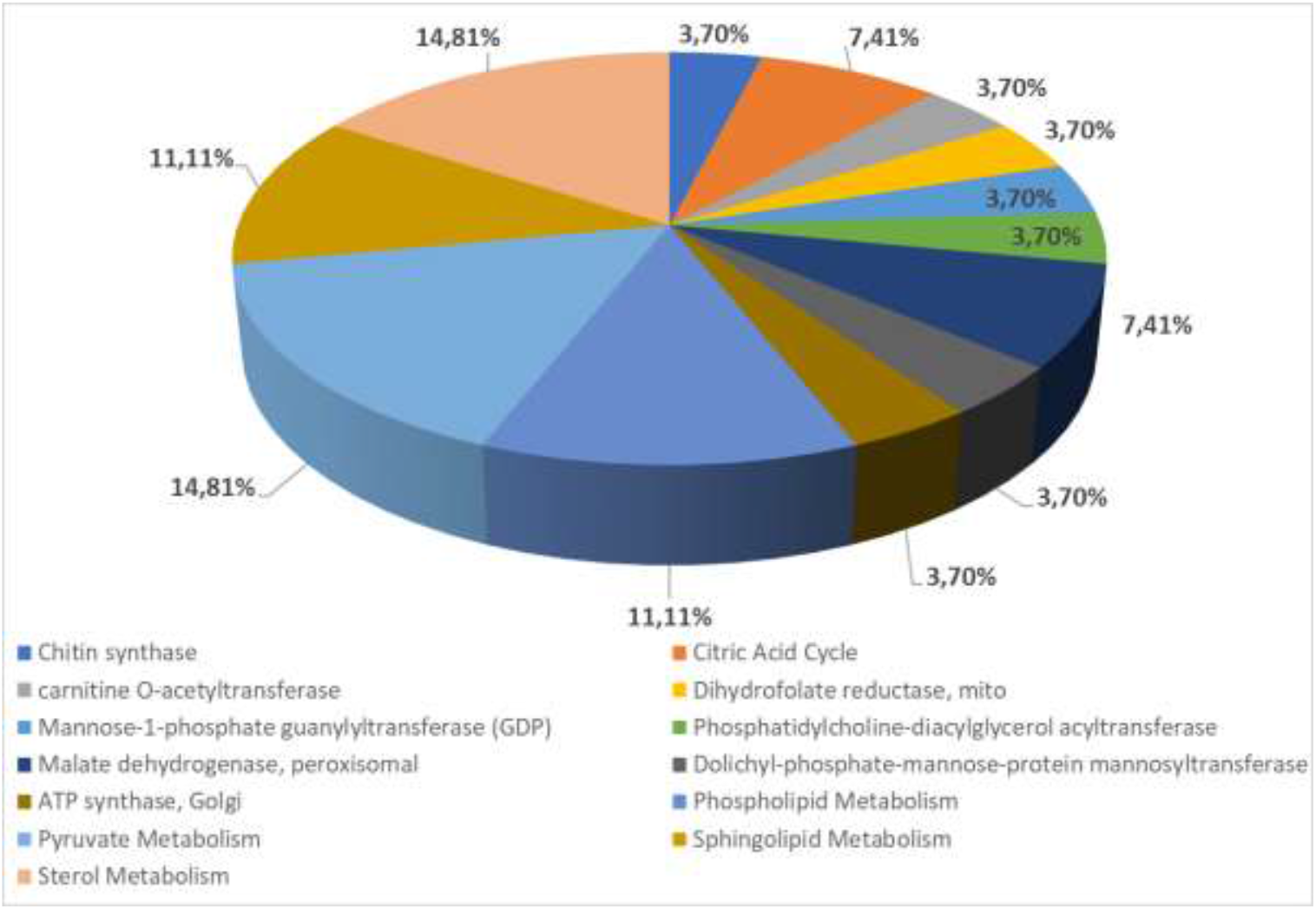
Downregulation/deletion suggestions for other metabolic pathways and compartments.

The algorithm suggests that these reactions and pathways have an indirect effect on heme production and must be detailed and analysed with additional omics data measurements during the cultivation process.

Downregulation of sterol metabolism might be related to the fact, that within the sterols desaturation reactions in yeasts - proteins contain one or two heme and Fe atoms in their active sites^52^, therefore maximising surplus heme production can be achieved by downregulation their consumption.

Summarising all described above we would like to point out the most promising up/down-regulation and deletion suggestions.

For upregulation, the best candidates are heme-BIOSYNTHESIS-II and PWY-5189, which include all heme biosynthesis pathways. Previous studies showed that HEM3 (HMBS reaction) is a rate-limiting step in *S. cerevisiae*^17^, but in *P pastoris* it is not. More, experimental data showed that *HEM1* (ALASm), *HEM2* (PPBNGS), *HEM3* (HMBS) and *HEM12* (FCLTm) upregulation leads to a 41 % increase in intracellular heme production and all HEBE biosynthesis reactions upregulation lead only to 1,5-fold increase.

In comparison with *S. cerevisiae*, the latest report shows 70-fold intracellular heme increase potential by upregulating heme biosynthesis pathways and deleting not essential carbon or nitrogen-consuming reactions like serine hydroxymethyltransferase (GHMT2r), heme oxygenase (biliverdin-producing) (heme-OXYGENASE-DECYCLIZING-RXN in MetaCyc^53^) and glycine cleavage system (GLYCLm). Heme-OXYGENASE-DECYCLIZING-RXN is not found in the iMT1026 GEM, thus hasn’t been calculated step-weighted factor, but in other organisms, the reaction converts heme to bilirubin, which would rapidly decrease intracellular heme concentration.

GLYCm reaction converts glycine and THF to 5,10-Methylenetetrahydrofolate and is used as a response to high concentrations of the glycine, thus diverting and decreasing it from heme production. GHMT2r is responsible for converting serine to glycine using thf as a co-factor. Our model suggests it as an upregulation candidate but published results showed that deletion will in total increase intracellular heme. This is explained because our algorithm does not include quantitative omics measurements which more precisely explain enzyme activity, available amount and indirect effects on other metabolic processes. Nevertheless, comparing within literature published data algorithm shows some inconsistencies, still, it shows feasible upregulation (33 reactions) (1.47 % of total reactions) and 67 downregulation/deletion (2.98 % of total) reaction suggestions results. Since GSM optimisations use methanol cultivation results, then regulation and deletion suggestions are based on *P. pastoris* methanol cultivation conditions data and parameters.

## Materials and methods

### Genome-scale metabolic model

#### Growth-coupled analysis

To analyse GEM for growth-coupled reactions and gene count manipulation impact on heme biosynthesis, we improved *P. pastoris* iMT1026 GEM, which consists of 1708 metabolites, 2243 reactions, 2 different biomass compositions for glycerol and methanol fermentation optimisations, has 9 different compartments and transport intercompartment for Cobra toolbox 3.0 compatibility^41^.

We developed an algorithm based on a previously published modelling approach for Lycopene production improvement for gene amplification targets^42^. The main concept of the framework is to select reaction amplification targets for improved product formation. The framework concept is to search for the candidate reaction, which flux changes have the most impact on target metabolite production under maximising biomass reaction flux. During analysis 3 different enzymatic reaction types were determined: flux-increasing reaction, flux-decreasing reaction and unaffected. In the end algorithm develops MS Excel file with exact results in specific tables (Supplementary materials 3), the content of each table is listed in Supplementary materials 2.

Before using the growth-coupled GEM modelling algorithm, there is a need to find the non-growth ATP maintenance cost. The model must be able to reach a steady state and produce ATP, which is necessary for maintenance processes. This is shown in Fig.3 where the biomass formation minimal flux value must be calculated and minimal ATP consumption must be maintained, otherwise, the model will not reach a steady state due to not enough ATP metabolic pool. After this value has been calculated, the algorithm finds the theoretical maximal biomass formation flux rate. When the minimal and maximal biomass flux rates are determined, the growth-coupled GEM modelling algorithm calculates the heme production maximal flux value by setting it as an objective function and maximising using the Flux Balance Analysis approach (FBA)^43^. In this stage, the growth-coupled GEM modelling algorithm has done initiation steps and is ready to find out each GEM reaction’s impact on product formation rate. In our case product formation is the heme production rate. The next step was to count heme production flux value increase step count (Fig. 3 black dots), which was divided into 5 equal steps to obtain the most promising optimisation results. In each step, the algorithm determines the objective reaction flux value change ratio against product flux increase value. The results are saved in a Supplementary Materials 3 Table S3.2, where Reactions ID from GSM and reactions names are saved in the “Reactions_ID” and “Reactions_name” columns and all 5 forced flux values of each reaction are saved in the “FBA_results_” columns. This table contains all found reactions with direct and indirect connections with heme production. More, we included subsystem and reactions stoichiometry information from GSM in the “Subsystem” and in the “Var10” columns to improve later data analysis steps.

**Figure 3.**
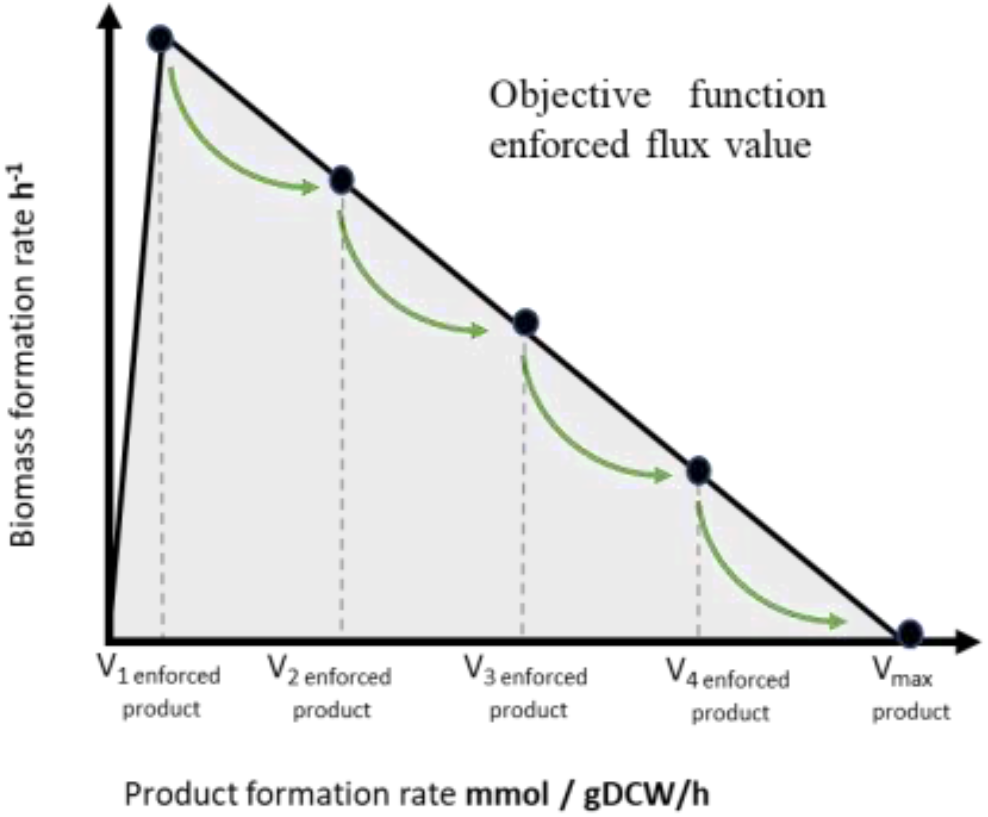
Growth-coupled GEM modelling algorithm concept.

After initiation and heme production flux value step count determination, the optimisation task was done. During optimisation, the last step for the heme transport reaction was set less than 10 % less of the maximal heme calculated value. This is important because in GEM reactions flux distribution is calculated by constraint-based flux analysis and, without additional omics data constraints implementation, the maximal flux rate constraint mostly is biologically irrelevant and biomass formation flux value becomes 0. In this case, there is no growth in the model and it is not possible to calculate growth-coupled reactions and genes manipulation impact on heme biosynthesis.

As a result, the algorithm determines 3 different GEM reaction types: flux-increasing reaction, flux-decreasing reaction and unaffected (Fig. 4). Flux-increasing reactions are considered for up-regulation, Flux-decreasing reactions are considered for down-regulation or deletion, for non-essential metabolites (the biomass building block chemicals or non-essential amino acids). There are also unaffected reactions, which do not impact heme production. The algorithm finds out all 3 types of reactions and saves them in the Supplementary materials 3 Table S3.2).

**Figure 4.**
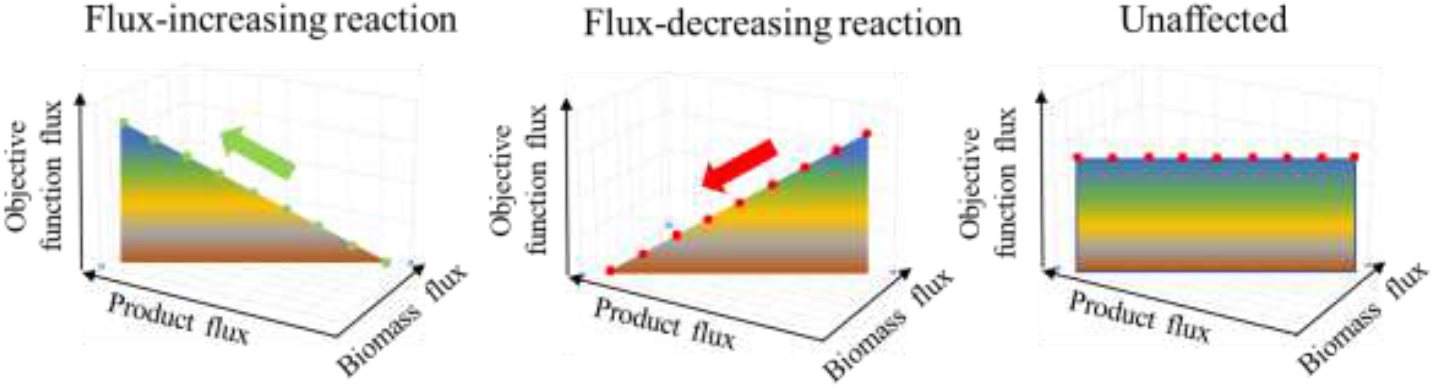
Reaction types identified by Growth-coupled GEM modelling algorithm.

After growth-coupled GEM modelling algorithm optimisations, we made excel additional reactions data filtering because not all irreversible reactions stoichiometry in GEM are in the same direction, which can lead to incorrect or incomplete optimisation results. Also, not all reactions had linearly changing flux values or even steady-state results, thus only reactions with all 5 results and steadily changing reaction flux values were chosen for the next analysis steps. As result, the algorithm generates 4 additional MS Excel data tables:

1. “Positive_contra_proportional”- reactions with flux value from left to right and are inversely proportionally to forced product (heme) changing fluxes;
2. “Negative_contra_proportional” - reactions with flux value from right to left and are inversely proportionally to forced product (heme) changing fluxes;
3. “Positive_directly_proportional” - reactions with flux value from left to right and are directly proportionally to forced product (heme) changing fluxes;
4. “Negative_directly_proportional” - reactions with flux value from right to left and are directly proportionally to forced product (heme) changing fluxes.

When all upregulation and downregulation/deletion reactions are filtered out and saved in separate data tables (Supplementary materials 3 Table S3.3-S3.6), then additional calculations are made. The algorithm finds and writes in the MS Excel file, the Boolean value if the reaction flux is left to right or right to left. This is necessary if the algorithm makes unpredicted inconsistencies, then manual filtering in the file is available.

### Step-weighted factor

To determine the most direct proportional or inversely proportional impact on objective product improved production, we introduced the step-weighted factor (SWF), which was calculated as (Formula 1):

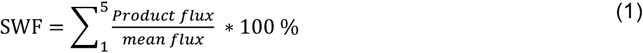

The weighted factor shows how much percentage each flux value changes versus mean value and are summed together. As not all reactions flux changes are in constant value (Supplementary materials 3 Table S3.2), then we summed all reaction changes and calculated the sum of percentages. The bigger number the better impact on the target product (increased heme production). Reactions with the largest step-weighted factor are the main candidates for reaction flux and gene regulation to increase heme production yields. Additionally, we calculated each reaction essentiality, which is reaction deletion essential for growth. For downregulation/deletion candidates only non-essential reactions were used (Supplementary Materials 3 Table S3.8).

### Candidate reaction analysis

After all, the reactions are sorted out, and the algorithm will filter the most potential ones, which upregulation could lead to a heme production increase. The second step is to find out which reactions must be downregulated/deleted to increase heme production. Deletion of the specific reaction in living cells can sometimes lead to lethality. To find out essential reactions in the filtered most potential reactions list we performed a single-reaction deletion analysis using CobraToolbox 3.0. In the end, we got reactions deletion list and genes deletion list suggestions for experimental genetic manipulation strain designing.

### Construction of the expression vector and selection of clones

An artificial gene encoding the soy legHemoglobin sequence (GenBank Acc. NP_001235248.2) was designed by GenScript and synthesized by BioCat GmbH (Heidelberg, Germany). The gene was cloned into the pPICZC vector (Invitrogen) using EcoRI and NotI restriction sites. The plasmid was linearized with PmeI and transformed in *Pichia pastoris* strain X-33 by electroporation. Mut+ transformants were produced on agarized YPD plates containing 800 μg/mL zeocin and analytically cultivated in flasks with rich BMMY medium for three days to select the best producer clone.

### Experimental conditions

Batch cultivation and feeding media solutions used in this study were prepared according to the “Pichia fermentation process guidelines” by Invitrogen Corporation^54^. 1.9 l of Basal Salts Medium: 26.7 ml L^−1^ H_3_PO_4_ 85 %, 0.93 g L^−1^ CaSO_4_, 18.2 g L^−1^ K_2_SO_4_, 14.9 g L^−1^ MgSO_4_·7H_2_O, 4.13 g L^−1^ KOH, 40.0 g L^−1^ glycerol and 4.35 ml L^−1^ PTM1 trace-element solution (0.02 g L^−1^ H_3_BO_4_, 5 ml L^−1^ H_2_SO_4_ 98 %, 6.0 g L^−1^ CuSO_4_·5H_2_O, 0.08 g L^−1^ NaI, 3.0 g L^−1^ MnSO_4_·H_2_O, 0.2 g L^−^1 Na_2_MoO_4_·2H_2_O, 0.5 g L^−1^ Ca_2_SO_4_·2H_2_O, 20.0 g L^−1^ ZnCl_2_, 65.0 g L^−1^ FeSO_4_·7H_2_O, 0.2 g L^−^1 biotin), were inoculated with 100 mL of inoculum grown in BMGY medium (10.0 g L^−1^ yeast extract, 20.0 g L^−1^ peptone, 100 mM potassium phosphate buffer, pH 6.0, 13.4 g L^−1^ yeast nitrogen base, 10.0 g L^−1^ glycerol, and 0.0004 g L^−1^ biotin) at 30 °C for 18–22 h in a shake flask at 250 RPM. Two feeding solutions were used - glycerol fed-batch solution (50 % glycerol, 12 mL L^−1^ PTM1) and methanol fed-batch solution (100 % methanol, 12 mL L^−1^ PTM1).

The bioreactor vessel was filled with distilled water and sterilized at 121 °C for 30 minutes, while the cultivation mediums and glycerol fed-batch solutions were autoclaved separately at the same conditions. The PTM1 and methanol fed-batch solutions were sterilized by filtration through a 0.2 μm filter.

The fermentations were conducted in a 5 L bench-top fermenter (Bioreactors.net, EDF-5.4/BIO-4, Latvia) with a working volume of 2-4 L. The pH was monitored using a calibrated pH sensor probe (Hamilton, EasyFerm Bio, Switzerland) and adjusted to 5.0 ± 0.1 with a 28 % NH_4_OH solution prior to starting the cultivation process and maintained at the set value during fermentation. The temperature was controlled at 30.0 ± 0.1 °C, using a temperature sensor and by adjusting the temperature in the vessel jacket. The dissolved oxygen (DO) level was measured using a DO probe (Hamilton, Oxyferm Bio, Switzerland) and kept above 30 ± 5 % by varying the stirrer speed (200-1000 RPM) (Cascade 1) or enriching the inlet air with pure O_2_ (Cascade 2). A constant air or air/oxygen mixture at a flow rate of 3.0 slpm was maintained throughout the process. A condenser was used to condense moisture from outlet gasses, and antifoam 204 (Sigma) was added when necessary to control excessive foam formation. Substrate feed solutions were pumped using a high-accuracy peristaltic pump (Longer-Pump, BT100–2J, China).

Process O_2_ and CO_2_ concentration were measured in the reactor exhaust gas using an O_2_/CO_2_ analyser (Bluesens, BlueInOneFerm, Germany). Culture turbidity was estimated using a turbidity probe (Optek, ASD19-EB-01, Germany) measuring the light absorption (transmission) within the 840– 910 nm wavelength range. Finally, a permittivity probe (Hamilton, Incyte, Switzerland) was used to estimate the viable cell concentration during cultivation.

The cultivations began with a glycerol batch phase. After 18-24 hours, the batch glycerol was depleted, and a glycerol fed-batch solution was fed into the reactor at a rate of 0.61 mL/min for 4 hours, or until an optical density of 100-120 was achieved. After a brief feeding pause of 10-30 minutes, to allow the cells to consume all of the residual glycerol, the feed substrate was switched to methanol, and the solution was fed into the reactor at a rate of 0.12 mL/min for 5 hours, followed by 0.24 mL/min for 2 hours, and finally 0.36 mL/min for the remainder of the cultivation.

### Analytical measurements

Cell growth was observed by offline measurements of the culture optical density (OD) at a wavelength of 590 nm (GRANAT, KFK-2, St. Petersburg, Russia). Wet cell weight (WCW) measurements were determined gravimetrically. Biomass samples were placed in pre-weighted Eppendorf® tubes and centrifuged at 15’500g for 3 minutes. Afterwards, the supernatant was discarded, the cells resuspended in distilled water and centrifuged once more. The liquid phase was discarded and the remaining wet cell biomass was weighed. Dry cell weight (DCW) was determined using a previously-determined empirical formula:

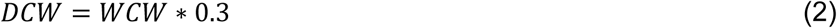

## Supporting information

https://github.com/BigDataInSilicoBiologyGroup/Pichia_pastoris_metabolic_modeling/tree/main/Supplementary_materials

## Data availability

All data generated or analysed during this study are included in this published article and supplementary materials. All supplementary materials are found in the link: https://github.com/BigDataInSilicoBiologyGroup/Pichia_pastoris_metabolic_modeling.

## Acknowledgements

This research was co-funded by European Regional Development Fund Project No. 1.1.1.1/21/A/044 “The development of an efficient pilot-scale leghemoglobine production technology, based on recombinant *Pichia pastoris* and *Kluyveromyces lactis* fed-batch fermentations. (BioHeme)”.

## Author contributions statement

A.P. and J.L. performed genome-scale metabolic model development, E.B. and A.K. constructed the expression vector and selected clones, E.B., Anastasija S., E.D., K.D. conceived the experiments, J.V. and Arturs S. conceived and coordinated the project. All authors reviewed the manuscript.

## Additional Information

The authors declare no competing interests.

## Notes

https://github.com/BigDataInSilicoBiologyGroup/Pichia_pastoris_metabolic_modeling

### Competing Interest Statement

The authors have declared no competing interest.

