## Supplementary material for "*Pichia pastoris* growth - coupled heme biosynthesis analysis using metabolic modelling": https://github.com/BigDataInSilicoBiologyGroup/Pichia_pastoris_metabolic_modeling/tree/main/Supplementary_materials

**Supplementary materials 1**

**CER, OUR and cell biomass estimation**

Carbon evolution rate (CER) and oxygen uptake rate (OUR) were estimated using the O_2_/CO_2_ analyzer and calculated using the following formulas:

$$CER= \frac{F_{air}*P}{V*R*T}*\left( \frac{1-O_{2}\%in-{CO}_{2}\%in}{1-O_{2}\%out-{CO}_{2}\%out}*{CO}_{2}\%out-{CO}_{2}\%in \right)$$

(1)

$$OUR= \frac{F_{air}*P}{V*R*T}*\left( O_{2}\%in-\frac{1-O_{2}\%in-{CO}_{2}\%in}{1-O_{2}\%out-{CO}_{2}\%out}*O_{2}\%out \right)$$

(2)

where, Fair – inlet air flow (L/h), P – normal pressure (1.0133 bar), V – bioreactor working volume (L), R – gas constant (8.314*10-2 bar L K-1 mol-1), T – process temperature (K), O_2_%in and O_2_%out – oxygen concentration in the inlet and outlet gas, respectively (%), CO_2_%in and CO_2_%out – carbon dioxide concentration in the inlet and outlet gas, respectively (%).

Cell biomass concentration was estimated using the Optek (turbidity) and Incyte (permittivity) sensor probes. Correlations between sensor signals and cell biomass were established using the experimental measurements as reference.

The Incyte sensor measures culture dielectric permittivity, and, thus, correlates only with the viable cell fraction in the bioreactor, which is preferable for metabolic modelling, as reactions can happen only in viable cells. As the experimentally measured cell concentrations include also the dead cell fraction, an empirical cell death rate coefficient (µd = 0.033 h-1) can be estimated using the sensor data. This coefficient can further be applied to the Optek sensor measurement, which normally takes into account also the dead cell fraction, to correlate the signal with only vegetative cells. Thus, two different viable cell estimation procedures were established and further used in metabolic model validation.

**Table S1.** Dry cell biomass (X_dry_), CER, OUR and specific growth rate estimations, based on Optek (turbidity) and Incyte (permittivity) sensor measurements at the end of each cultivation phase.

|  | **X_dry_ (g/L)** | **CER (mmol g^-1^ h^-1^)** | **OUR (mmol g^-1^ h^-1^)** | **µ (h^-1^)** |
| --- | --- | --- | --- | --- |
| *Optek sensor measurement* | | | | |
| Glycerol | 62.5 | 1.61 | 2.17 | 0.19 |
| Methanol | 89.0 | 1.13 | 1.76 | 0.03 |
| *Incyte sensor measurement* | | | | |
| Glycerol | 59.2 | 1.70 | 2.29 | 0.19 |
| Methanol | 110.85 | 0.91 | 1.41 | 0.04 |

**Supplementary materials 2**

**Optimisation “Optimisation_result_Final” MS Excel file Content**

1) “Table S3.1. All_in_one” – all *P. pastoris* iMT1026 genome-scale metabolic model reactions.
2) “Table S3.2. Dependant_reactions” – all reactions which have an impact on ferroheme *b production.*
3) “Table S3.3. Pos_contra_proportional”- reactions with flux value from left to right and are inversely proportionally to forced product (heme) changing fluxes;
4) “Table S3.4. Neg_contra_proportional” - reactions with flux value from right to left and are inversely proportionally to forced product (heme) changing fluxes;
5) “Table S3.5. Pos_directly_proportional” - reactions with flux value from left to right and are directly proportionally to forced product (heme) changing fluxes;
6) “Table S3.6. Neg_directly_proportional” - reactions with flux value from right to left and are directly proportionally to forced product (heme) changing fluxes.
7) “Table S3.7. Directly_proportional” – All reactions which are directly proportional. Excluded all transport reactions, reactions with less than 5 flux values, and inconsistent flux values.
8) “Table S3.8. Inversely proportional” – All reactions which are directly proportional. Excluded all transport reactions, reactions with less than 5 flux values, and inconsistent flux values.
9) “Table S3.9. Inversly_reactions” – Inversely proportional reaction group list which is displayed in Figure 2.
10) “Table S3.10. Inversly_amino” - Inversely proportional reaction group list which is displayed in Figure 1.
